# The Relationship between Stability of Interpersonal Coordination and Inter-Brain EEG Synchronization during Anti-phase Tapping

**DOI:** 10.1101/2021.04.20.440627

**Authors:** Yuto Kurihara, Toru Takahashi, Rieko Osu

## Abstract

Inter-brain synchronization is enhanced when individuals perform rhythmic interpersonal coordination tasks, such as playing instruments in music ensembles. Experimentally, synchronization has been shown to correlate with the performance of joint tapping tasks. However, it is unclear whether inter-brain synchronization is related to the stability of interpersonal coordination represented as the standard deviation of relative phase (SDRP). In this study, we simultaneously recorded electroencephalograms of two paired individuals during anti-phase tapping in three speed conditions: slow (reference inter-tap interval [ITI]: 0.5 s), fast (reference ITI: 0.25 s), and free (preferred ITI). We calculated the inter-brain synchronization within six regions of interest: frontal, central, left/right temporal, parietal, and occipital regions. We found that synchronization of the central-temporal regions was positively correlated with SDRP in the theta and alpha bands, while synchronization of the frontal-frontal and frontal-central was positively correlated with SDRP in the beta band. These results demonstrate that inter-brain synchronization occurs only when task requirements are high, and that it increases with the instability of the coordination. This may be explained by the stronger mutual prediction required in unstable coordination than that in stable coordination, which increases inter-brain synchronization.

## Introduction

People interact during group dancing and music ensembles by coordinating their actions swiftly and accurately [1]. These widespread social activities involve temporally precise interpersonal synchronization based on the information exchanged via multiple sensory modalities [2]. Furthermore, these activities require that individuals coordinate stably to exhibit their performance [3][4]. Previous studies have examined interpersonal coordination using simple joint tapping tasks, such as in-phase or anti-phase tapping between two individuals [4][5][6].

Interpersonal coordination patterns can be represented by a relative phase (RP) that captures the angular differences between two oscillators [7][8][9][10]. The standard deviation of relative phase (SDRP) represents the instability of the coordination patterns. Two patterns of interpersonal coordination have been examined, in-phase (RP=0°) and anti-phase (RP=180°) modes [11]. In-phase coordination is more stable than anti-phase coordination [7][8]. In particular, the anti-phase interpersonal coordination becomes increasingly unstable (increase in SDRP) as the movement frequency increases, eventually breaking down and transiting to in-phase coordination (generally called phase transition) [7][12]. For instance, Schmidt et al. observed that, when two participants coordinated leg movements with one another, the SDRP for the anti-phase mode was larger than that for the in-phase mode, and transition from the anti-phase to in-phase coordination was noted when the frequency of leg movement increased [7].

Recently, to elucidate the neural mechanisms of interpersonal coordination, hyperscanning has been used to examine the synchronization of two or more brains (inter-brain synchronization) [13][14] during a variety of interaction tasks from simple joint tapping tasks [15][16] to complex natural tasks, such as conversations [17]. Previous research has also demonstrated the relationship between inter-brain synchronization of electroencephalogram (EEG) signals and behavioral performance, indicating the degree of achievement in an interpersonal coordination task [18]. Dumas et al. found increased alpha-band inter-brain EEG synchronization in the right centroparietal area when the imitator’s movements were synchronized with the model’s movements [19]. Yun et al. found an increased inter-brain synchronization of theta and beta frequency band EEGs during an implicit fingertip synchrony task after leader-follower movement training [20]. Furthermore, Kawasaki et al. suggested that good performance pairs of visually guided alternate tapping showed higher inter-brain EEG synchronization in the alpha frequency (12 Hz) than poor performance pairs [16]. These previous hyperscanning studies have focused on behavioral performance representing the degree of accomplishment of the task required by the experimenter. However, none have examined the relationship between inter-brain synchronization and the stability of interpersonal coordination.

The relationship between stability and brain activity has been studied during bimanual coordination [21]. For instance, using functional magnetic resonance imaging (fMRI), Lindenberg et al. found that premotor and supplemental motor cortices were activated when instability in bimanual coordination of the index finger increased [22]. In addition, transcranial magnetic stimulation of these regions caused instability of bimanual coordination. Aramaki et al. reported prefrontal, premotor, and parietal activations in relation to the phase transition of bimanual finger-tapping movements [23]. Aramaki et al. demonstrated that the initiation-related transient brain activity in the putamen is useful for predicting the future stability of periodic bimanual movements [24]. The putamen shows greater activation during anti-phase movements than during in-phase movements [25][26]. In this way, the stability of bimanual coordination influences activities of various regions (frontal, motor, parietal regions, and putamen) of the brain. These brain regions may be associated with instability of interpersonal coordination during the anti-phase tapping.

This study aimed to elucidate the relationship between the stability of interpersonal coordination (SDRP) and inter-brain EEG synchronization during anti-phase tapping, which is more unstable than in-phase tapping, especially when the tapping speed is increased [7]. In addition, the anti-phase mode can avoid the spurious inter-brain EEG synchronization caused by similar behaviors across individuals during in-phase tapping [27]. Therefore, we hypothesized that if stability reflects performance, inter-brain EEG synchronization would be high in a stable condition. On the other hand, during the bimanual coordination task, the brain is activated when the instability is high [22]. Therefore, our other hypothesis was that the inter-brain synchronization would increase as the instability increases. To evaluate these hypotheses, we assessed SDRP and inter-brain synchronization during anti-phase finger tapping in slow, fast, and free speed conditions.

## Results

### Behavioral measurements during anti-phase tapping

We calculated the inter-tap interval (ITI) [5][6][28] by subtracting the tap onset time of one participant from the subsequent tap onset time of the other to see whether participants performed anti-phase tapping to the reference ITI (Figure 1). Figures 2A and 2B depict ensemble-averaged time series of ITI and RP and their standard deviation (SD). In order to confirm that averaged ITI and RP time series were stationary processes, we performed Dickey-Fuller tests [29]. The averaged ITI and RP time series were confirmed as stationary processes in the slow, fast, and free conditions.

**Figure 1:**
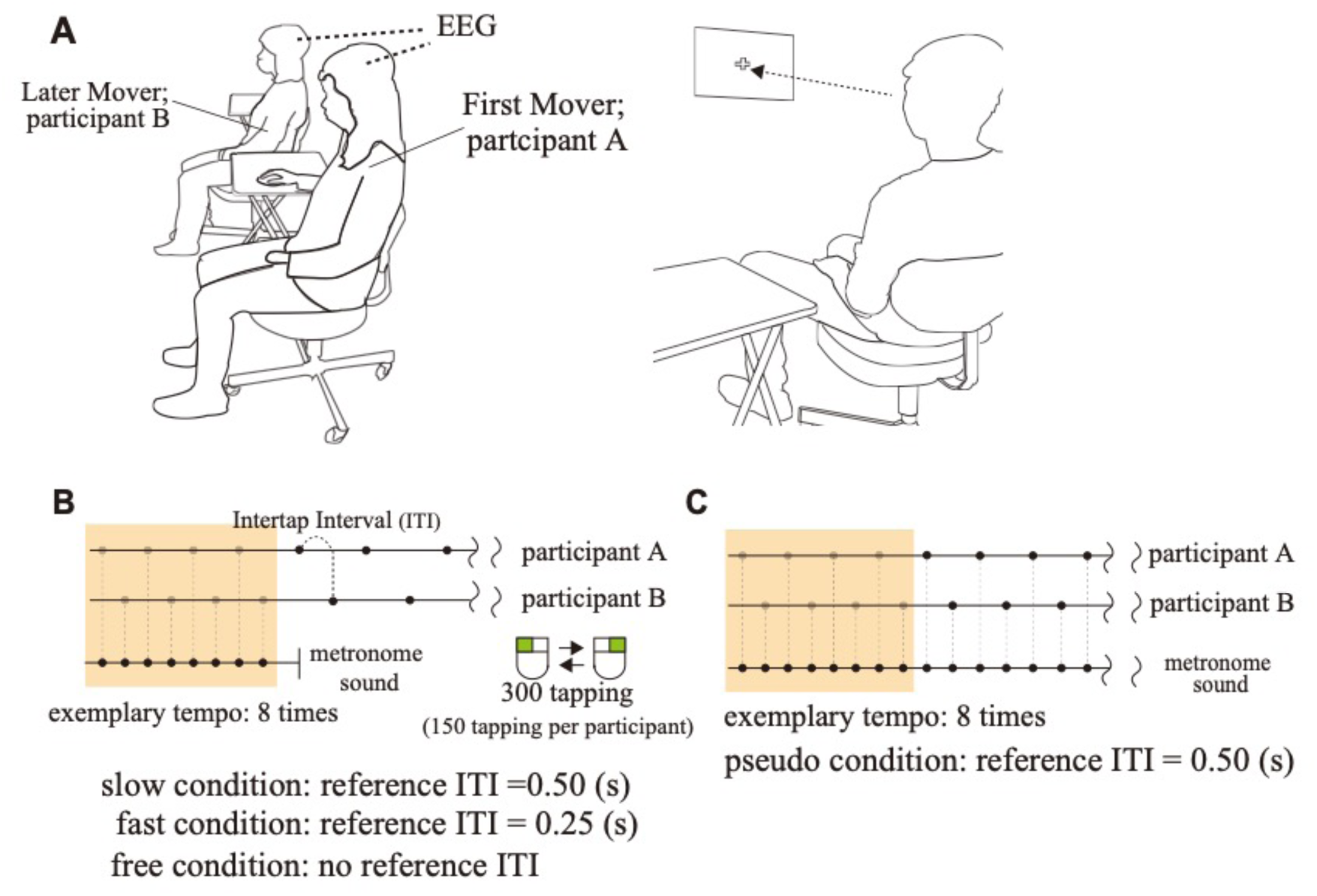
Experimental setting and the procedure of anti-phase tapping tasks. (A) We conducted hyperscanning using two wireless EEG systems. Each participant gazed at a fixation point in front of him/her during anti-phase tapping. (B) The anti-phase tapping tasks consisted of slow, fast, and free speed conditions. In the slow and fast conditions, the participants first listened to exemplary sounds (reference ITI) of a slow (2 Hz) and a fast (4 Hz) frequency. After the participants listened to the sound, they started to perform anti-phase tapping with the same frequency as that of the reference ITI. In the free condition, there was no reference sound. Thus, the participants performed the tapping with preferred frequency in the free condition. (C) The experiment also involved pseudo conditions in which participants performed anti-phase tapping matched to metronome sounds of 2 Hz.

**Figure 2:**
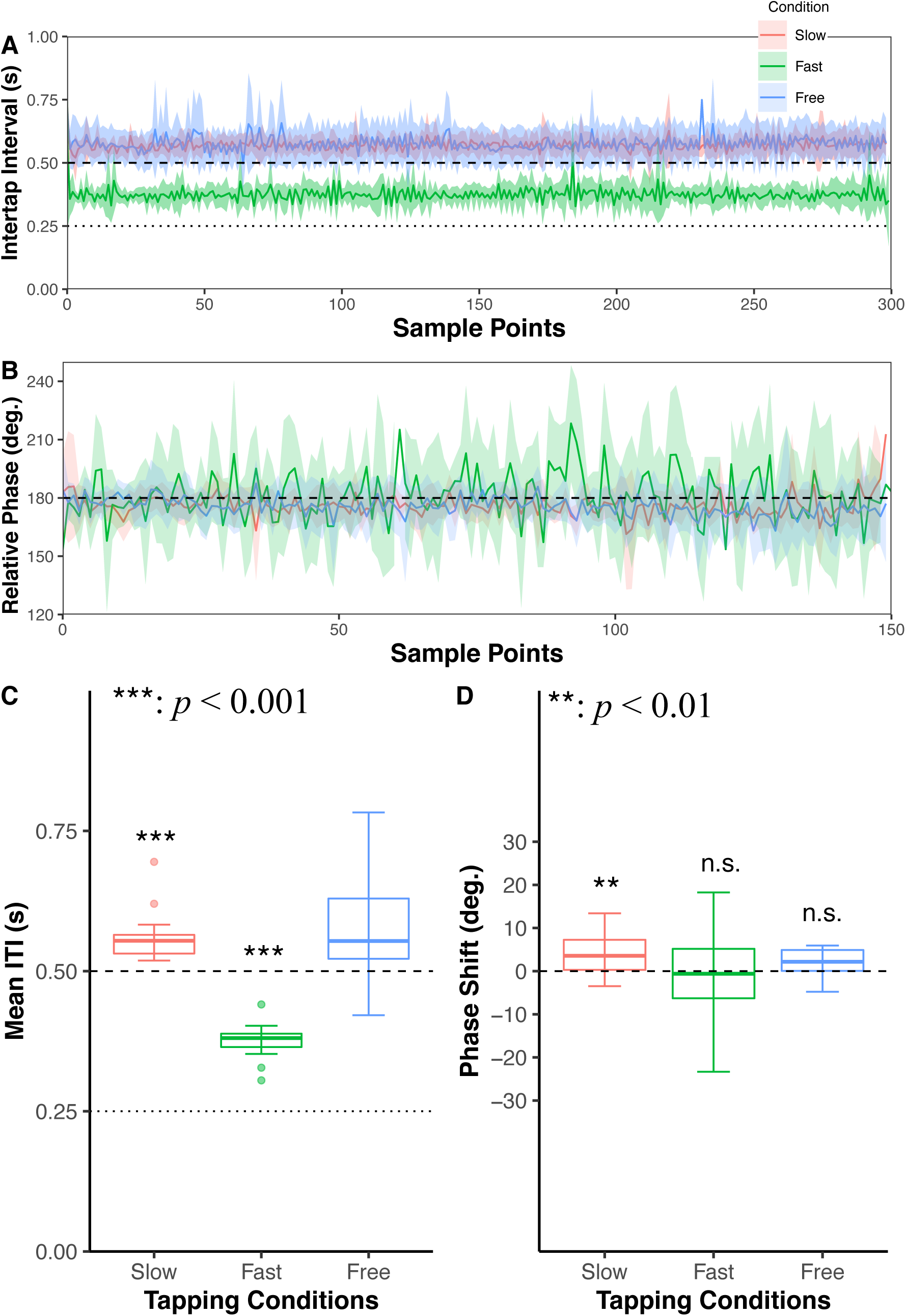
The behavioral analysis of anti-phase tapping in slow, fast and free tapping condition. (A) The curves show the ensemble average of inter-tap interval (ITI) time series across participants. The light color areas indicate the standard error of ITI. The dashed line indicates the reference ITI in the slow condition. The dotted line indicates the reference ITI in the fast condition. (B) The curves show the ensemble average of the relative phase (RP) time series across participants. The light color areas indicate the standard error of RP. The dashed line indicates the anti-phase angle. (C) The box plots show the median and interquartile range (IQR) of the mean inter-tap intervals (mean ITI). In the free condition, the one-sample t-test was not conducted with the mean ITI because of no reference ITI. (D) The box plots show the median and IQR of the phase shift of tapping. \**p*<0.05, \*\**p*<0.01, and \*\*\**p*<0.001.

We calculated the average of ITI (mean ITI), phase shift (shift from 180° computed by subtracting the RP from 180°), and SDRP (standard deviation of RP). First, a one-sample t-test revealed that the mean ITI was significantly different from the reference ITI in the slow and fast conditions (slow, *t*12=4.9579, *p<*0.001; fast, *t*12=13.406, *p<*0.001) (Figure 2C). The average and SD of the mean ITI in the free condition was 0.577±0.10 (s). Thus, we confirmed that the mean ITI was longer than the reference ITI in slow and fast conditions. In addition, a one-sample t-test was conducted for the phase shift to confirm the shifting from reference RP (180°) in slow, fast, and free conditions (Figure 2D). The phase shift was significantly different from the anti-phase degree (180°) in the slow condition (*t*12=−3.0567, *p*=0.009). On the other hand, the phase shift was not significantly different from the anti-phase degree (180°) in fast and free conditions (fast, *t*12=−1.139, *p*=0.276; free, *t*12=1.3664, *p*=0.1969). In the slow condition, the RP was slightly higher than the anti-phase tapping (mean=4.1037°, SD=4.841°). These results indicate that the participants could typically perform accurate anti-phase tapping.

The one-way repeated measures analysis of variance (ANOVA) was conducted for SDRP to compare slow, fast, and free conditions (Figure 3). The assumption of sphericity was examined by Mauchly’s test. The degree of freedom was not corrected because the assumption of sphericity was not violated (*p=*0.064). ANOVA with tapping conditions revealed the main effects of tapping conditions on SDRP (*F*2,24*=*8.928, *p=*0.001*, ηp2=*0.427; Figure 3). Post-hoc paired t-tests using a Holm correction confirmed that the SDRP of the fast condition was larger than those of slow and free conditions (slow vs. fast, *t12=*−3.734, *padj=*0.003*, d=*−1.036; fast vs. free, *t12=*3.580, *padj=*0.003*, d=*0.993; and slow vs. free, *t*13=*−*0.154, *padj=*0.879, *d=−*0.043). Thus, participants performed the anti-phase tapping with high variability in the fast condition.

**Figure 3:**
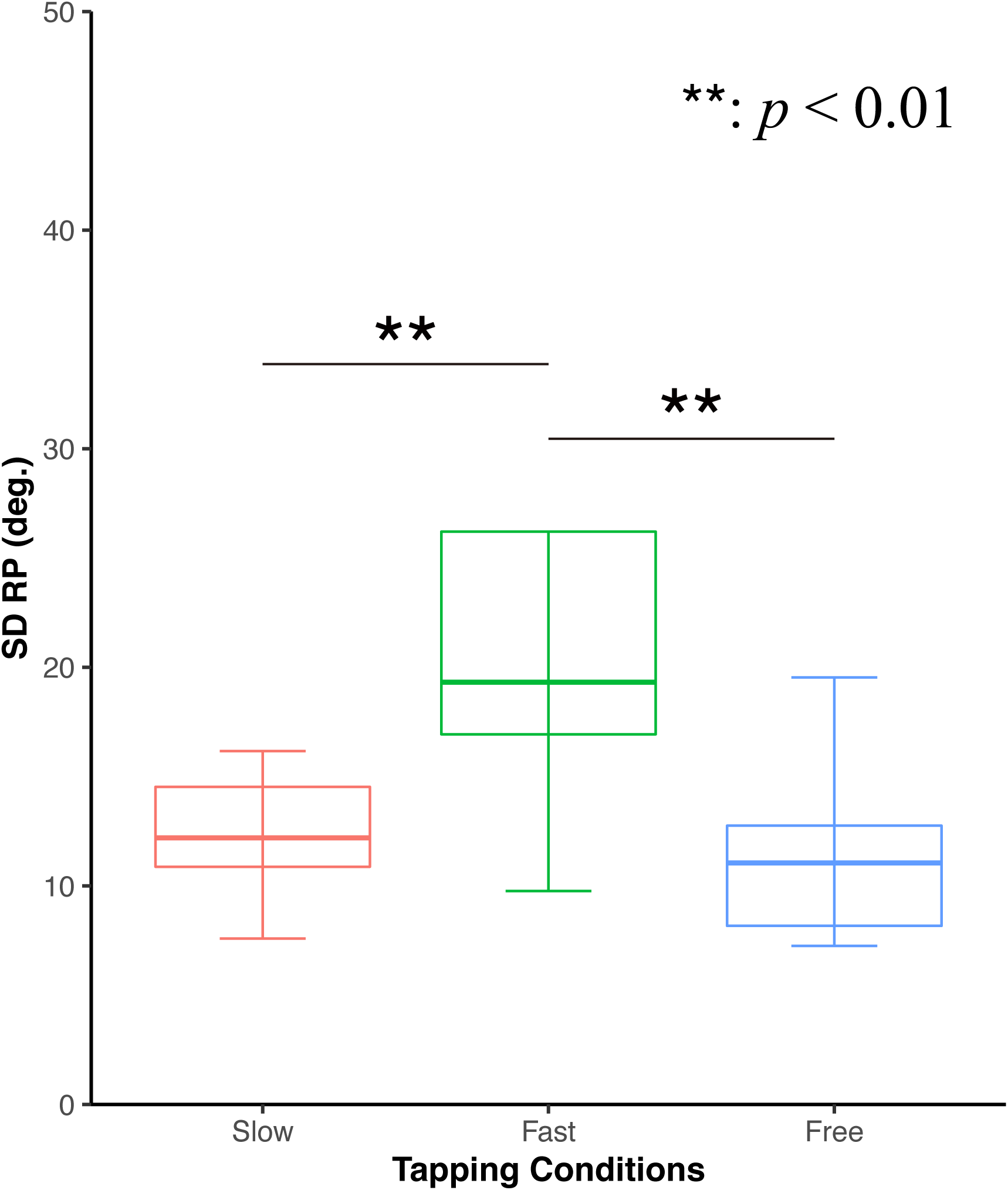
The difference of SDRP among slow, fast, and free tapping conditions. The SDRP in fast conditions was significantly larger than those in slow and free conditions.

### Comparison between original and surrogate EEG synchronizations

The inter-brain EEG synchronization was evaluated using CCorr [30][31] for each electrode pair between two participants in theta, alpha, and beta frequency bands. Because each participant had 29 electrodes, we computed the CCorr of 841 possible combinations of electrode pairs. We categorized the 29 electrodes into six regions of interest (ROIs): frontal (Fp1, Fp2, AF3, AF4, F7, F8, F3, F4, and Fz), central (FC5, FC6, C3, C4, and Cz), left temporal (T7, CP5, and P7), right temporal (T8, CP6, and P8), parietal (P3, P4, Pz, PO7, PO8, PO3, and PO4), and occipital (O1 and O2) [17][32][33] (Figure 4), resulting in 21 ROI pairs (see methods for detail). Then, we averaged the CCorrs of electrode pairs within each ROI pair.

**Figure 4:**
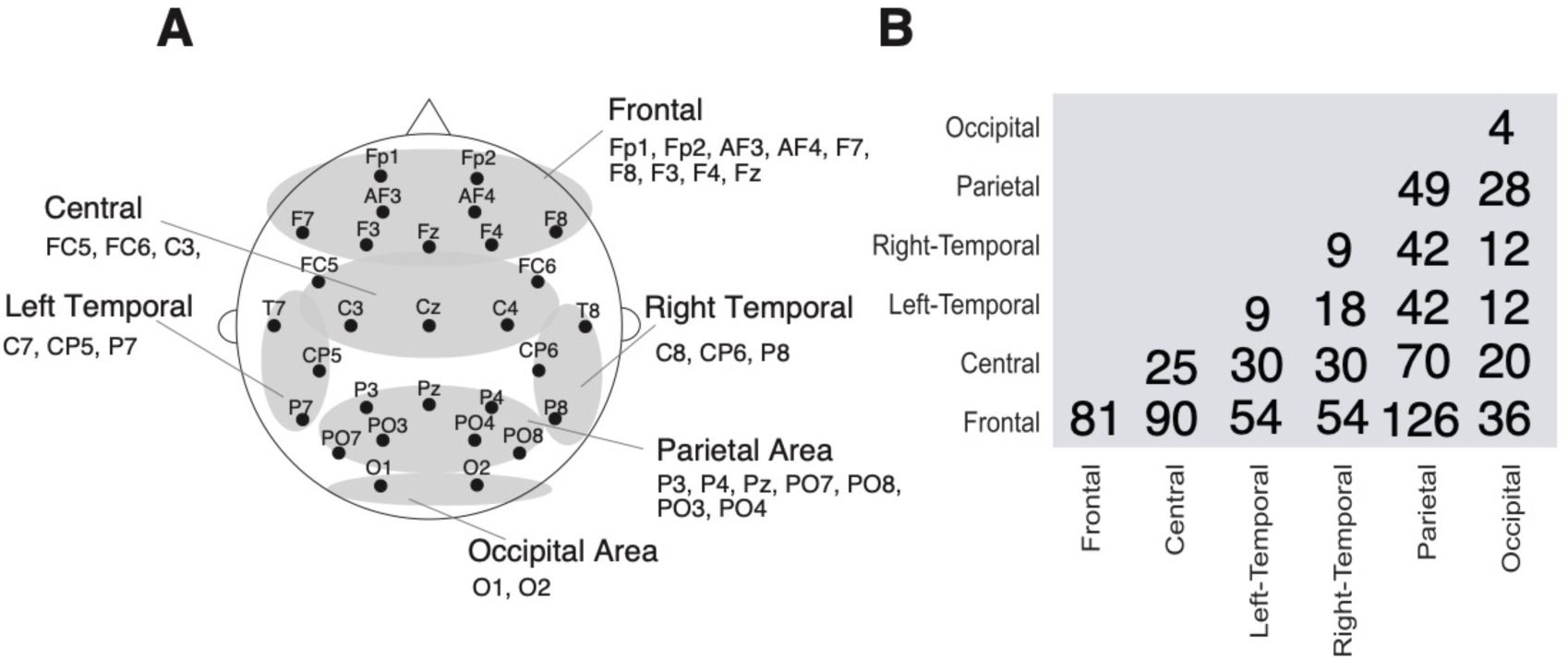
Six regions of interest and the number of channel pairs included in each ROI pairs. (A)The electrodes were classified into six regions of interest: frontal (Fp1, Fp2, AF3, AF4, F7, F8, F3, F4, and Fz), central (FC5, FC6, C3, C4, and Cz), left temporal (T7, CP5, and P7), right temporal (T8, CP6, and P8), parietal (P3, P4, Pz, PO7, PO8, PO3, and PO4), and occipital (O1, and O2) areas. (B) The matrix shows the number of inter-brain channel pairs between which CCorr were computed for each ROI pairs.

To examine whether the observed EEG synchronization was obtained by chance, we created surrogate data from the original data and compared the difference in CCorr between original and surrogate data for 21 ROI pairs and conducted Welch’s t-test (one-tail) with false discovery rate (FDR) correction (number of comparisons, 21 ROI pairs) for each experimental condition and frequency band (Figure 5). Among CCorrs of 21 ROI pairs, we found three significantly larger CCorrs and 15 marginally significantly larger CCorrs than surrogate CCorrs in the theta bands in the fast condition. In the fast condition, we also found 10 significantly larger and eight marginally significantly larger CCorrs than surrogate in the alpha bands, 15 significantly larger and four marginally significant CCorr than surrogate in the beta bands. There were not significantly larger CCorrs in the other condition or other frequency bands. Consequently, CCorrs of theta, alpha, and beta frequency bands were significantly larger in the fast condition.

**Figure 5:**
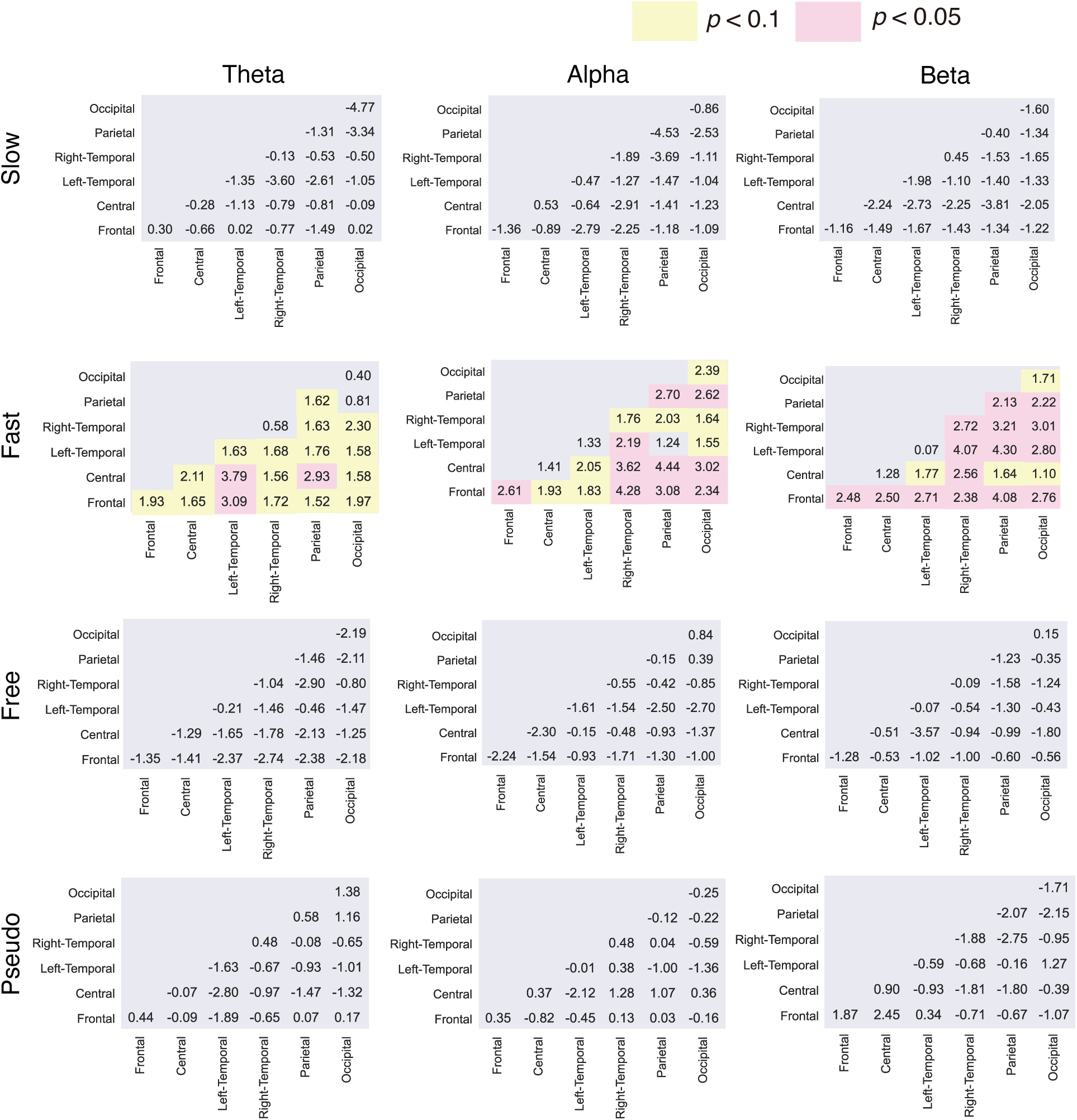
Comparison between original and surrogate CCorr in theta, alpha, and beta frequency bands. The matrices contain the t-values of Welch’s t-test comparing original and surrogate CCorrs in each pair of region of interest in slow, fast, free, and pseudo conditions. ROI pairs that had significantly larger CCorrs between participants during anti-phase tapping compared to surrogate are indicated by red in the corresponding cells with *padj* <0.05 and by yellow in the corresponding cells with *padj* <0.1.

### Relationship between inter-brain EEG synchronization and the stability of anti-phase tapping

We calculated the Spearman correlation between SDRP and average CCorrs of the ROI pairs that were significantly or marginally significantly larger than surrogates in the fast condition (see above). In theta frequency bands, the SDRP was marginally significantly correlated with the CCorr between the left temporal and central ROIs (*ρ=*0.621, *padj=*0.071; Figure 6A). The SDRP also significantly positively correlated with the right temporal-central CCorr in the alpha frequency band (*ρ=*0.775, *padj=*0.021; Figure 6B). Furthermore, there were marginally significant positive correlations between the SDRP and CCorrs in frontal-frontal and frontal-central ROI pairs in the beta bands (frontal-frontal, *ρ=*0.742, *padj=*0.056; frontal-central, *ρ=*0.775, *padj=*0.077; Figure 6C).

**Figure 6:**
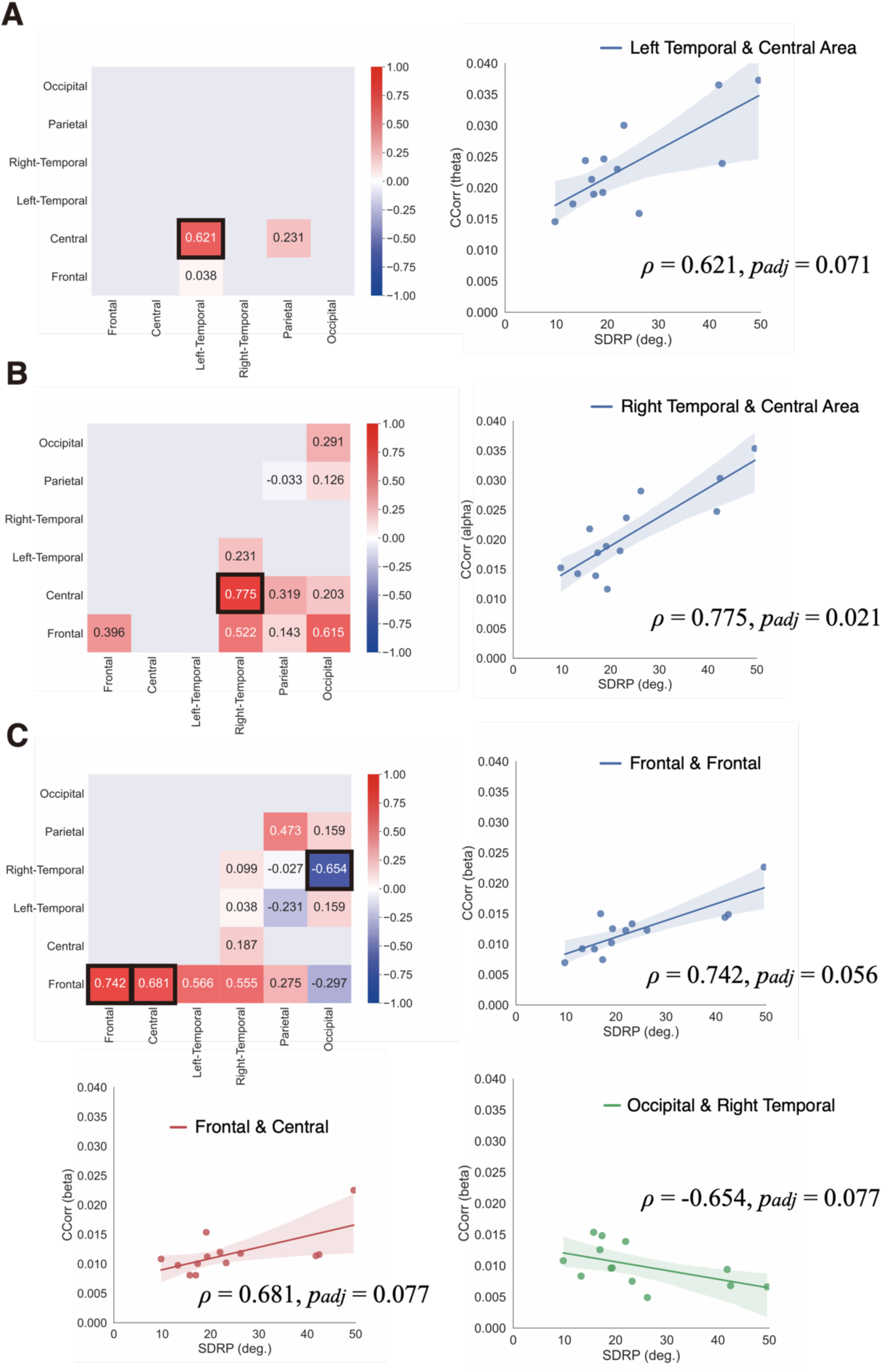
Relationship between inter-brain electroencephalogram synchronization and the instability of anti-phase tapping in the fast condition (A) The heatmap indicates Spearman correlations between SDRP and CCorr in the theta frequency band. The scatter plot shows a marginally significant positive correlation between SDRP and CCorr between the left-temporal and the central region. (B) The heatmap indicates Spearman correlations between SDRP and CCorr in the alpha frequency band. The scatter plot shows a significant positive correlation between SDRP and CCorr between the right-temporal and the central region. (C) The heatmap indicates Spearman correlations between SDRP and CCorr in the beta frequency band. The upper right scatter plot (blue color) shows a significant positive correlation between SDRP and CCorr between the frontal-frontal region. The lower left scatter plot (red color) shows a marginally significant positive correlation between SDRP and CCorr between the frontal and the central regions. The lower left scatter plot (green color) shows the marginally significant negative correlation between SDRP and CCorr between the occipital and the right-temporal regions.

Thus, in these regions, the inter-brain synchronization increased when the tapping was less stable. There was a marginally significant negative correlation in occipital-right temporal regions in the beta bands, i.e., inter-brain synchronization was larger when the tapping was more stable (*ρ=*−0.654, *padj=*0.077; Figure 6C).

## Discussion

This study examined the relationship between the behavioral stability of interpersonal coordination and inter-brain EEG synchronization. We observed significantly high inter-brain EEG synchronization in multiple ROIs in the theta, alpha, and beta bands during anti-phase tapping with fast speed as compared to the surrogate. Among these ROIs, we found a positive correlation between tapping instability (SDRP) and inter-brain CCorr between the left temporal region of one participant and the central regions of the other paired participant in the theta frequency bands. The SDRP positively correlated with the inter-brain CCorr between right temporal and central regions in the alpha frequency band. The SDRP positively correlated with the CCorr between the frontal and the frontal regions, as well as between the frontal and the central regions in the beta frequency band. In contrast, the SDRP was negatively correlated with the inter-brain CCorr between the occipital and the central regions in the beta frequency bands.

Overall, these results reveal a relationship between tapping instability and inter-brain EEG synchronization in the frontal, central, and left/right temporal regions with the fast speed condition, while there was a relationship between tapping stability and inter-brain synchronization in the occipital-right temporal regions in the beta frequency band with the fast speed condition.

In this study, we focused on the SDRP of anti-phase tapping. Previous studies exhibited more SDRP for anti-phase than for in-phase coordination and spontaneously transitioned from anti-phase to in-phase coordination as the coordination was unstable [7][12]. Thus, the SDRP of anti-phase tapping can be regarded as a behavioral performance index which implies that participants maintain anti-phase tapping. Our results show that with worsened anti-phase tapping performance (high SDRP), inter-brain synchronization increases in the frontal, central, and temporal regions in theta, alpha, and beta bands. These results appear to be in opposition to those of Kawasaki et al., who proposed that inter-brain synchronization is related to good performance [16]. However, because the two indices differ between the studies it is not possible to directly compare them. Kawasaki et al. regarded good performance when the interval between two consecutive taps was <50 ms, and assessed the binary expression (i.e., good/poor), while SDRP was a continuous variable. There is also a possibility of different degrees of difficulty, as Kawasaki et al. conducted the tapping task at a preferred speed (∼0.50 s) that was equivalent to the slow and free conditions in our study.

The increased inter-brain EEG synchronization observed in unstable tapping may be explained by the mutual prediction theory, which indicates that a participant tries to predict the behavior of their partner. According to the mutual prediction theory, each brain has systems to control behavior, and at the same time, systems to predict the behavior of others [34]. Cui et al. found larger inter-brain synchronization in the right superior frontal cortices during a cooperation task that required prediction of the partner’s behavior than during a competition task [35]. Cooperation tasks requires a high degree of mutual prediction [34] for coordination with one another, whereas competition tasks do not require mutual prediction. In addition, Liu et al. reported an increased inter-brain synchronization in the dorsomedial prefrontal cortex (dmPFC) during the cooperative condition in a Jenga game [36] that required a high mutual prediction. Other studies have observed an increased inter-brain synchronization during cooperation tasks [37][38][39][40]. Individuals appear to rely on prediction in order to maintain a certain tempo during joint tapping because individuals depend on an anticipatory mechanism to achieve good synchrony [41][42][43]. Performing fast anti-phase tapping would require each participant to accurately predict the behavior of their partner to keep a constant tempo because anti-phase tapping tends to become unstable (high SDRP) at a fast tempo, which similarly occurs during bimanual coordination [12][44]. In contrast, it is easy for participants to keep anti-phase tapping at a slow tempo (slow, free, and pseudo conditions), which therefore requires less prediction between partners. The state of low stability of interpersonal coordination requires cooperation with each other. Therefore, there is the possibility that the observed increase in inter-brain synchronization is associated with mutual prediction, which needs to sustain the stability of interpersonal coordination. Thus, our results may be explained by the mutual prediction theory.

We assumed that inter-brain synchronization increases as the instability increase because, the brain was found to be activated when the instability of bimanual coordination was high. Bimanual coordination of the anti-phase mode requires brain activity to maintain the stable coordination pattern [26]. In hyperscanning during interpersonal coordination, participants also require inter-brain synchronization to predict and maintain mutually anti-phase tapping. Therefore, the increased brain activity of unstable bimanual coordination is consistent with inter-brain synchronization of unstable interpersonal coordination in that it requires the brain activity to maintain stable coordination.

We observed significant inter-brain EEG synchronization in theta (4-7 Hz), alpha (8-12 Hz), and beta (13-30 Hz) frequency bands. Our study found more inter-brain EEG synchronization in the beta band than theta and alpha bands in the fast condition (Figure 6). It is not yet clear which frequency band is significantly related to interpersonal coordination [45]. Yun et al. observed a significant increase in the inter-brain EEG synchronization in theta and beta frequency bands during unintentional body movements [20]. Kawasaki et al. showed that the inter-brain EEG synchronization in the alpha frequency (about 12 Hz) was higher in good tapping pairs than that in poor tapping pairs [16]. In the interpersonal coordination task of finger tapping, stimulating the left M1 of the pairs on beta rhythm (20 Hz) using transcranial alternating current stimulation (tACS) enhanced tapping performance [46]. In our tapping experiments, the sound was feedback. Previous EEG and MEG studies indicate that beta-band oscillation represents auditory rhythm in frontal, temporal regions [47][48]. However, in our study, there were no temporal regions related to inter-brain EEG synchronization in the beta frequency band, but frontal and central regions. Thus, the feedback sounds are not the only cause for why inter-brain EEG synchronization occurs. From ours and previous studies, inter-brain EEG synchronization during interpersonal coordination is considered to be related to the beta frequency band. However, no studies have conclusively shown which frequency band is essential to interpersonal coordination.

Our results reveal the possibility that the frontal-frontal inter-brain synchronization might be related to the instability of interpersonal coordination. The identified area seems to match several previous EEG hyperscanning studies. Yun et al. observed increased inter-brain synchronization during their finger keeping task in the inferior frontal gyrus, anterior cingulate cortex, and ventromedial prefrontal cortex (vmPFC) identified by sLORETA (standardized low resolution brain electromagnetic tomography) after leader-follower movement training [20]. Cui et al. found that the inter-brain synchronization between signals generated by the right superior frontal cortices increased during cooperation tasks [35]. Furthermore, from dual brain experiments in mice, inter-brain synchronization in the prefrontal cortex was found to be related to social interaction [49]. Our result of the inter-brain synchronization in the frontal area is consistent with previous hyperscanning studies. In addition, we found inter-brain EEG synchronization between signals generated by central and temporal regions in the theta and alpha frequency bands. In our task, participants inputted the sounds and outputted the tapping behaviors. Temporal regions respond to sounds [50] and central regions respond to tapping coordination [51]. Therefore, there may be inter-brain synchronization between the central and temporal regions. To identify the area more accurately, EEG source estimation using individual MRI structure is preferable, which is one of our future directions.

Our study has the following limitations. First, the small sample size limited the statistical power of the study. Future study with large sample size will clarify the relationship between interpersonal coordination and inter-brain synchronization in more detail. Second, the two paired participants each heard the same tapping sound, which is unavoidable for anti-phase tapping task. Indeed, there is the possibility that two brains showed the similar neural activity simply because they receive the same auditory input. Third, the classification of EEG cannels into ROIs used in this study is not necessarily a standardized method. Future study should conduct EEG source estimation using individual MRI structures with more EEG channels. Fourth, although the CCorr is considered to be more robust against pseudo inter-brain synchronization than any other indices [30], it is unclear whether the CCorr best represents inter-brain interaction. Future study must compare different indices of inter-brain synchronization.

In this study, we found a correlation between the instability of interpersonal coordination and inter-brain synchronization. However, the causality between the instability and inter-brain synchronization remains unclear. The modulation of inter-brain synchronization using tACS [53] would reveal the causality between interpersonal coordination and inter-brain synchronization.

In summary, our findings support the hypothesis that inter-brain EEG synchronization is related to the stability of interpersonal coordination. However, our results reveal that the correlation is negative, i.e., inter-brain synchronization increased as the behavior became more unstable in the frontal, central, and temporal regions in theta, alpha, and beta bands. It is possible that high inter-brain synchronization promotes cooperation to maintain the stability of interpersonal coordination. The results may be explained by the mutual prediction theory.

## Methods

### Ethics

The experimental procedures were approved by the Ethical Review Committee of Waseda University. All experiments were performed in accordance with relevant guidelines and regulations. All participants provided written informed consent.

### Participants

Nineteen pairs of right-handed participants (8 male pairs, 11 female pairs; mean age=22.5 years, SD=4.3) were enrolled. Two male pairs and four female pairs were excluded because of recording artifacts: two female pairs because of device trouble and the others because of EEG artifacts. Thus, 13 pairs (eight acquaintance and five stranger pairs) were included. The acquaintances knew each other, while the strangers had never seen each other prior to the experiments.

### Experimental settings and behavioral task

The participants in each pair were seated side-by-side (Figure 1A). They were asked to gaze at a fixation point during the tapping task to avoid possible synchronization resulting from mutual glances. The distance between the participants was approximately 70 cm. The participants were asked to perform anti-phase tapping tasks using two computer mice. These computer mice were placed on two tables (40 × 50 cm). The participant who tapped first used the mouse’s left click with his/her right index finger.

The other participant used the right click of the mouse with his/her right index finger. Tapping produced sound feedback (sound frequency, 440 Hz). Each participant listened to the sound produced by his/her own tap and his/her partner’s tap through earphones.

The experiment included four tapping conditions that consisted of different tapping frequencies: slow condition (reference ITI=0.50 s), fast condition (reference ITI=0.25 s), free condition (an ITI preferred by the pair), and pseudo condition (tapping according to 0.50 s metronome sounds). In the slow and fast conditions, the participants first listened to the exemplary frequency (Figure 1B). After the first eight sounds were transmitted, the metronome was switched off, and the pairs started tapping at a frequency as close as possible to the memorized reference frequency. In the free condition, there was no reference frequency; the participants tapped at a preferred frequency. In the pseudo condition, after eight sounds of 2 Hz, the participants continued tapping to the 2 Hz metronome. There was no sound feedback to both participants in the pseudo condition. A trial in each condition was completed after 300 taps. Therefore, the slow, fast, free, and pseudo conditions had different durations for one trial (slow: mean, 175.24 s, SD, 14.35 s; fast: mean, 116.96 s, SD, 10.08 s; free: mean, 178.94 s, SD, 29.44 s; pseudo: mean, 160.28 s, SD, 0.294 s). All conditions consisted of only one trial. The order of the first and second mover who started tapping was fixed across all conditions.

### Behavioral analysis

We calculated the ITI by subtracting the tap onset time from the adjacent tap onset time. ITI was defined using tapping times as follows:

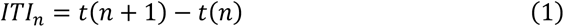

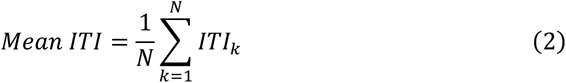

where t(n) denotes the n-th tap timing, N is the number of tapping, mean ITI indicates the average of ITI, and SDITI indicates the standard deviation of ITI. We calculated the RP of tapping using a circular measure to confirm whether tapping was anti-phase (RP=180°)[9][46]. RP was defined using tapping times:

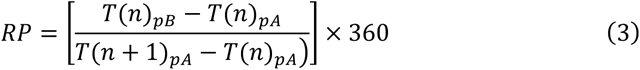

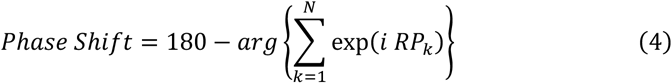

where *T*(*n*)*_pA_*(and *T*(*n*)*_pB_* denote the tapping time (second) of participants A and B, respectively. The RP ranged from 0° to 360°. The phase shift indicates the average RP of shifting from 180° using circular statistics. The *i* shows the complex number and the arg {·} shows the argument of complex. We calculated the circular standard deviation of RP (SDRP) as follows:

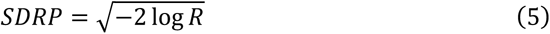

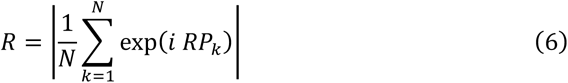

where SDRP represents the instability of anti-phase tapping. R is the resultant length of mean RP vector.

### EEG hyperscanning data recording and analyses

The brain activities of each pair were simultaneously recorded by two EEG systems, each with 29 active scalp electrodes (Quick-30; Cognionics, San Diego, CA, USA) in accordance with the placement of the international 10/20 system: Fp1/Fp2, AF3/AF4, F3/F4, F7/F8, FC5/FC6, C3/C4, T7/T8, CP5/CP6, P3/P4, P7/P8, PO3/PO4, PO7/PO8, O1/O2, Fz, Cz, and Pz. The sampling rate was 500 Hz. Reference electrodes were placed on the left ear lobe.

EEG preprocessing was conducted using Python 3.8.5 with MNE-Python (0.20.7). The EEG data were down-sampled to 250 Hz. Next, the EEG data were filtered with a band-pass ranging from 1-45 Hz to remove artifacts. Each EEG channel was visually inspected for bad channels. To reduce or eliminate artifacts (electrooculogram, muscle noise, sweating, and movement), we conducted an independent component analysis (ICA) on the EEG. After ICA, we removed independent components derived from artifacts. Bad channels were then interpolated using spherical spline interpolation. The EEG data were band-pass filtered to extract the theta (4-7 Hz), alpha (8-12 Hz), and beta (13-30 Hz) frequency bands.

The inter-brain EEG synchronization was evaluated using CCorr [30][31] for each electrode pair between two participants. CCorr was directly parallel to the Pearson product-moment correlation coefficient for the circular data.

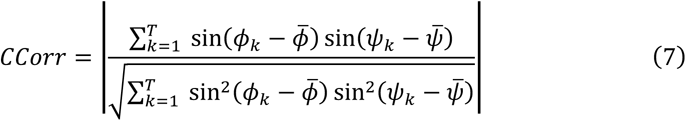

where *ϕ_k_* and *ψ_k_* are the kth EEG phases of participants A and B, extracted using the Hilbert transform from theta, alpha, and beta frequency band passed signals, respectively. *ϕ̄* and *ψ̄* are the mean directions for the phase of participant A and B, respectively. *k* is the time point of down sampled EEG (250 Hz), and T is the number of EEG time points of participants A and B of the corresponding trial. The CCorr values ranged from 0 to +1. CCorr was adopted in this study because it is less susceptible to detecting spurious hyper-connections of type one error connectivity [30]. The CCorr is much more robust to coincidental synchronization because it measures the phase variance of oscillations. Burgress suggested that the phase locking value (PLV) cannot measure the circular co-variance, and the PLV might introduce pseudo-connectivity from the results of the hyper-connectivity simulation [30].

To detect the region of inter-brain synchronization related to the stability of anti-phase tapping, we created the six ROIs for the EEG electrodes (Figure 4). We calculated the average CCorrs for each ROI pair as follows: 1) we calculated the inter-brain CCorrs over every possible combination of inter-brain EEG electrodes (total of 841); 2) we averaged the CCorrs included in each ROI pair. For example, for the occipital-occipital ROI pair, CCorrs of four channel pairs, i.e., O1-O1, O1-O2, O2-O1, and O2-O2, were averaged. In addition, the CCorrs were averaged across symmetrical ROI pairs (e.g., for the right temporal-occipital ROI pair, CCorrs of the 12 cannel pairs, i.e., O1-T8, T8-O1, O1-CP6, CP6-O1, O1-P8, P8-O1, O2-T8, T8-O2, O2-CP6, CP6-O2, O2-P8, and P8-O2 were averaged) because of no distinction between the two participants of a pair, resulting in 21 ROIs pairs (see Fig. 4 for the number of channel pairs averaged in each ROI pair).

### Surrogate dataset

To elucidate if these inter-brain EEG synchronization results were obtained by chance, we created surrogate data and compared the original EEG synchronization (original CCorr) with surrogate EEG synchronization (surrogate CCorr). This surrogate CCorr was created as follows: 1) we created 1000 ms windows of one participant’s EEG time series in each pair of channels in slow, fast, free, and pseudo conditions and shuffled the order of time series (1000 ms window) across four conditions; 2) we extracted the 156 s (average time for tapping in four tapping conditions) windows from the shuffled EEG data; 3) we calculated the surrogate CCorr for each pair of EEG channels; and 4) we averaged the surrogate CCorr within each ROI pair. We conducted these four steps 50 times and created 50 average surrogate CCorrs for each ROI pair. The same surrogate CCorrs were used across the all tapping conditions to compare with original CCorrs. These steps were applied for theta, alpha, and beta frequency band, respectively.

### Statistical analyses

To confirm whether the ITI and RP time series were stationary processes, we conducted the Dickey-Fuller test [29]. If the *p*-value was <0.05, the time series represented the stationary process. Next, the one-sample t-test was conducted with the mean ITI in comparison to the reference ITI in slow and fast conditions. We also conducted the one-sample t-test for a phase shift to assess the shifting from 0° in slow, fast, and free conditions. The pseudo condition was excluded from the analysis of behavior because participants did not interact with each other in the pseudo condition.

One-way ANOVA was performed for SDRP to confirm the difference in the tapping stability among tapping conditions. The assumption of sphericity was examined by Mauchly’s test. When the *p*-value of Mauchly’s test was <0.05, the Greenhouse-Geisser correction was applied. After confirming the significance of the main effect of tapping conditions, post-hoc paired t-tests with Holm correction were used to examine significant differences among tapping conditions. These analyses were conducted using JASP (https://jasp-stats.org/).

To compare the differences between original and surrogate CCorrs in each ROI, we applied Welch’s t-test (one-tail) in slow, fast, free, and pseudo conditions in theta, alpha, and beta frequency bands. We corrected for multiple comparisons using FDR correction (number of comparisons, 21 ROI pairs). In subsequent analyses, we focused on the average CCorrs of ROI pairs which were significantly higher than the surrogate CCorrs. To confirm the relationship between inter-brain synchronization and the stability of interpersonal coordination, we calculated the Spearman correlation between these CCorrs and SDRP of anti-phase tapping in each ROI. These correlations’ p-values were corrected by FDR.

## Acknowledgments

We thank S.Okazaki for his experimental programming of tapping task. This research was supported by Grants-in-Aid for Scientific Research (KAKENHI) on Innovative Areas from the Ministry of Education, Culture, Sports, Science and Technology (MEXT), Japan, “Oscillogy” (Grand No. 18H04954) and “Construction of the Face-Body Studies in Transcultural Conditions” (Grand No. 20H04586).

## Author contributions statement

Y.K. and R.O. designed the experiments and prepared the manuscript. Y.K. collected and analyzed the data. T.T. and R.O. provided important suggestions on the data analysis and the manuscript. Y.K., T.T. and R.O. wrote the paper.

## Data availability

The datasets generated and analyzed during the current study are available from the corresponding author on reasonable request.

